# Central norepinephrine transmission is required for stress-induced repetitive behavior in two rodent models of obsessive-compulsive disorder

**DOI:** 10.1101/855205

**Authors:** Daniel Lustberg, Alexa Iannitelli, Rachel P. Tillage, Molly Pruitt, L. Cameron Liles, David Weinshenker

**Author notes:** Address correspondence to: David Weinshenker, Ph.D., Department of Human Genetics, Emory University School of Medicine, 615 Michael St., Whitehead 301, Atlanta, GA 30322.

## Abstract

**Rationale:** Obsessive-compulsive disorder (OCD) is characterized by repetitive behaviors exacerbated by stress. Many OCD patients do not respond to available pharmacotherapies, but neurosurgical ablation of the anterior cingulate cortex (ACC) can provide symptomatic relief. Although the ACC receives noradrenergic innervation and expresses adrenergic receptors (ARs), the involvement of norepinephrine (NE) in OCD has not been investigated.

**Objective:** To determine the effects of genetic or pharmacological disruption of NE neurotransmission on marble burying (MB) and nestlet shredding (NS) in two animal models of OCD.

**Methods:** We assessed NE-deficient (*Dbh -/-*) mice and NE-competent (*Dbh +/-*) controls in MB and NS tasks. We also measured the effects of anti-adrenergic drugs on NS and MB in control mice and the effects of pharmacological restoration of central NE in *Dbh -/-* mice. Finally, we compared c-fos induction in the locus coeruleus (LC) and ACC of *Dbh -/-* and control mice following both tasks.

**Results:** *Dbh -/-* mice virtually lacked MB and NS behaviors seen in control mice but did not differ in the elevated zero maze (EZM) model of general anxiety-like behavior. Pharmacological restoration of central NE synthesis in *Dbh -/-* mice completely rescued NS behavior, while NS and MB were suppressed in control mice by anti-adrenergic drugs. Expression of c-fos in the ACC was attenuated in *Dbh -/-* mice after MB and NS.

**Conclusion:** These findings support a role for NE transmission to the ACC in the expression of stress-induced compulsive behaviors and suggest further evaluation of anti-adrenergic drugs for OCD is warranted.

## Introduction

In patients with obsessive-compulsive disorder (OCD), intrusive thoughts drive repetitive and compulsive behaviors, among them checking, counting, and excessive hand washing. These obsessions are emotionally aversive and disproportionately command attentional resources, until the ritual that interrupts the obsession is completed (Graybiel and Rauch 2000; Pigott et al. 1994). OCD is highly comorbid with other psychiatric disorders, such as attention deficit hyperactivity disorder (ADHD), anxiety, and Tourette’s syndrome (Abramovitch et al. 2015; Carter et al. 2004; Eapen and Robertson 2000; Lombroso et al. 2008). In OCD and other compulsive disorders, symptom severity increases during periods of psychological stress (Adams et al. 2018; Lin et al. 2007; Rosso et al. 2012). The neural circuitry and neurochemistry that drive compulsive behaviors have not been clearly delineated (Micallef and Blin 2001; Stein 2000), but converging evidence from neuroimaging, neurosurgical, and neurobiological studies implicates a corticolimbic interface structure called the anterior cingulate cortex (ACC) in the pathophysiology of OCD (Alarcon et al. 1994; Brennan et al. 2013; Fitzgerald et al. 2005; McGovern and Sheth 2017).

Data from human functional imaging studies have frequently identified ACC hyperactivity in OCD patients (De Ridder et al. 2017; Fitzgerald et al. 2005; Riffkin et al. 2005; Van Laere et al. 2006). Both functional and structural abnormalities in the ACC of patients with OCD are correlated with disturbances in emotional reactivity, attention, and cognitive control (Brennan et al. 2015; McGovern and Sheth 2017; Pauls et al. 2014). The ACC receives glutamatergic and neuromodulatory input from a distributed network of forebrain and hindbrain structures and is well positioned to regulate attention, emotions, motivation, and action (Margulies et al. 2007; Yücel et al. 2003). For instance, the ACC sends glutamatergic projections to the striatum to initiate voluntary behavior, and ACC projections to the orbitofrontal and medial prefrontal cortex support stimulus evaluation, decision-making, and behavioral flexibility under conditions of uncertainty or change (Riffkin et al. 2005; Stevens et al. 2011). Findings from human and animal studies also demonstrate a functional role for the ACC in the response to environmental novelty (Krebs et al. 2013; Montag-Sallaz et al. 1999; Struthers et al. 2005; Tulving et al. 1994; Weible et al. 2009). In addition, neurons in the ACC may encode the aversive emotional component of pain sensation (Navratilova et al. 2015; Rainville et al. 1997; Shackman et al. 2011), and multiple studies have reported that hyperactivity of the ACC is associated with other aversive affective states (Hayes and Northoff 2011; Johansen and Fields 2004; Stevens et al. 2011).

Evidence from the neurosurgical literature suggests that ∼40% of antidepressant-resistant OCD patients can achieve lasting remission of symptoms by undergoing a surgical procedure called a cingulotomy (Brown et al. 2016; Kim et al. 2003; McGovern and Sheth 2017), in which the ACC is lesioned bilaterally with gamma radiation (Cosgrove and Rauch 2003; Fodstad et al. 1982). Despite the efficacy of this intervention for improving the quality of life in some treatment-resistant OCD patients, invasive surgical ablation of an important brain region is an aggressive and irreversible treatment strategy that should only be considered as a last resort after pharmacological interventions have failed. Indeed, cingulotomy is associated with significant long-term cognitive and emotional side effects, including emotional apathy and deficits in attention (Cohen et al. 1999a; Cohen et al. 1999b; Janer and Pardo 1991).

Currently, the only drugs approved for the management of OCD are serotonergic antidepressants (tricyclics and selective serotonin reuptake inhibitors; SSRIs), which are ineffective for about half of patients and unfortunately associated with adverse side effects that limit compliance (Dominguez 1992; Mavissakalian et al. 1983; Pizarro et al. 2014; van Balkom et al. 1994). Notably, very high antidepressant doses are required to improve OCD symptoms, and the latency for antidepressants to become effective is even longer for OCD patients than for patients with anxiety disorders or major depression (Dougherty et al. 2004; Fineberg and Gale 2005; Pittenger and Bloch 2014).

Because of the modest efficacy of SSRIs in alleviating OCD symptoms, most preclinical research on OCD pathophysiology has focused on the serotonin (5-HT) system and, to a lesser extent, the dopaminergic and glutamatergic systems (Billett et al. 1998; Goodman et al. 1990; Pittenger et al. 2011; Pittenger et al. 2006; Zohar et al. 2000). The development of more effective and tolerable pharmacotherapies has been impeded by an incomplete understanding of the neurobiological basis of OCD and other compulsive disorders (Langen et al. 2011; Micallef and Blin 2001; Ting and Feng 2011). Thus, there is an urgent need to develop and characterize better preclinical models of OCD to unravel the neurochemistry and neurocircuitry that drive compulsive behaviors.

Some advances in understanding the neurobiology of OCD have come from genetic and behavioral models of the disease, which have various degrees of face, construct, and predictive validity (Ahmari et al. 2015; Fineberg et al. 2011; Joel 2006; Ting and Feng 2011; Witkin 2008). The nestlet shredding (NS) and marble burying (MB) tasks are two rodent models of repetitive, compulsive behaviors that may be useful for testing novel pharmacotherapies for OCD (Angoa-Pérez et al. 2013; Li et al. 2006; Wolmarans et al. 2016). In these tasks, repetitive behaviors (digging for MB; shredding for NS) are elicited by cage-change stress and do not habituate even after repeated daily exposures to marbles or nestlets (Thomas et al. 2009; Witkin 2008).

Multiple studies have found that routine laboratory animal husbandry practices increase serum corticosterone levels and other physiological measures of stress (Balcombe et al. 2004; Okano et al. 2005; Rasmussen et al. 2011; Schultz 1972). Transferring mice to a clean cage or gently handling them with gloves, although seemingly innocuous, are associated with marked changes in affective behavior when animals are tested on the same day (Rasmussen et al. 2011). Sympathetic activation and endocrine stress responses to these procedures also do not habituate over a 15-day period (Balcombe et al. 2004).

Digging and nest building are normal rodent behaviors that are augmented by stress (Deacon 2006; Kedia and Chattarji 2014; Schultz 1972; Umathe et al. 2008). The fact that these behaviors do not habituate lends face validity to both tasks and distinguishes them from canonical models of novelty-induced anxiety (Handley 1991; Witkin 2008; Wolmarans et al. 2016). The construct validities of MB and NS are supported by studies using mice with genetic 5-HT depletion (*Tph2-/-*). These mice display exaggerated MB and NS behaviors (Angoa-Pérez et al. 2013; Angoa-Pérez et al. 2012; Kane et al. 2012). MB and NS also show predictive validity in pharmacological experiments; SSRIs and tricyclic antidepressants reduce these repetitive behaviors (Jimenez-Gomez et al. 2011; Li et al. 2006).

The central norepinephrine (NE) system mediates stress responses, attention, arousal, emotional state, and behavioral flexibility (Aston-Jones et al. 2007; Sara 2009), all of which are disrupted in patients with OCD (Adams et al. 2018; Cocchi et al. 2012; Gehring et al. 2000; Kalanthroff et al. 2016; Spitznagel and Suhr 2002). The 5-HT and NE neuromodulatory systems regulate one another (Kim et al. 2004; O’Leary et al. 2007; Pasquier et al. 1977; Pudovkina et al. 2002; Segal 1979) and have overlapping terminal fields in forebrain regions, such as the ACC, where 5-HT exerts an inhibitory influence (Czyrak et al. 2003; Hajós et al. 2003; Tanaka and North 1993) and NE exerts an excitatory influence (Berridge et al. 1993; Gompf et al. 2010; Marzo et al. 2014; Stone et al. 2006). Moreover, reciprocal excitatory communication between the noradrenergic locus coeruleus (LC) and the ACC is required to support sustained arousal during exposure to unfamiliar environments (Gompf et al. 2010), suggesting that the LC-ACC axis is a critical substrate of arousal and attention in response to contextual change (Aston-Jones et al. 1999; Vankov et al. 1995).

Although studies on the subject are limited, there is some compelling evidence for abnormalities in central NE function and adrenergic receptor (AR) sensitivity in patients with compulsive disorders. Elevated plasma levels of NE metabolites (Siever et al. 1983), altered neuroendocrine responses to adrenergic drug challenges (Brambilla et al. 1997; Hollander et al. 1991; Siever et al. 1983), and genetic polymorphisms in the NE-metabolizing enzyme *COMT* (Karayiorgou et al. 1999; Pooley et al. 2007; Schindler et al. 2000) have been reported in OCD patients, but the role of the central NE system in OCD pathophysiology in humans has not been investigated thoroughly. Similarly, the effects of serotonergic drugs on NS and MB behavior have been extensively documented, but the effects of drugs targeting the NE system have only been described in a handful of studies (Li et al. 2006; Millan et al. 2000; Sugimoto et al. 2007; Young et al. 2006).

Here we determined the effects of genetic or pharmacological disruption of central NE signaling on OCD-like behaviors in the NS and MB tasks using NE-deficient (*Dbh -/-*) mice and their NE-competent (*Dbh +/-*) counterparts (Thomas et al. 1995). To provide a contrast to canonical anxiety-like behavior, we also tested performance in the elevated zero maze (EZM). Finally, we assessed the effects of genetic NE deficiency on c-fos induction in the LC and ACC as a measure of task-specific neuronal activity during NS and MB tasks.

## Materials and methods

### Subjects

*Dbh* -/- mice were maintained on a mixed 129/SvEv and C57BL/6J background, as previously described (Thomas et al. 1998; Thomas et al. 1995). Pregnant *Dbh* +/- dams were given drinking water that contained the βAR agonist isoproterenol and α1AR agonist phenylephrine (20 µg/ml each) + vitamin C (2 mg/ml) from E9.5–E14.5, and L-3,4-dihydroxyphenylserine (DOPS; 2 mg/ml + vitamin C 2 mg/ml) from E14.5-parturition to prevent the embryonic lethality associated with the homozygous *Dbh* deficiency (Mitchell et al. 2008; Thomas et al. 1995). *Dbh* -/- mice are readily identified by their ptosis phenotype, and genotypes were subsequently confirmed by PCR. *Dbh* +/- littermates were used as controls because their behavior and catecholamine levels are indistinguishable from wild-type (*Dbh +/+*) mice (Marino et al. 2005; Mitchell et al. 2006; Thomas et al. 1998; Thomas et al. 1995).

Male and female mice 3–8 months old were used in all experiments. Because no sex differences were reported in the literature or observed in pilot experiments, male and female mice of the same *Dbh* genotype were pooled. All animal procedures and protocols were approved by the Emory University Animal Care and Use Committee in accordance with the National Institutes of Health guidelines for the care and use of laboratory animals. Mice were maintained on a 12 h light/12 h dark cycle with *ad libitum* access to food and water. Behavioral testing was conducted during the light cycle in a quiet room in which the animals were housed to minimize the stress of cage transport on test days.

### Drugs

The following drugs were used for behavioral pharmacology experiments: the non-selective β-adrenergic receptor (βAR) antagonist DL-propranolol hydrochloride (Sigma-Aldrich, St. Louis, MO), the α1AR antagonist prazosin hydrochloride (Sigma-Aldrich), the α2AR agonists guanfacine hydrochloride (Sigma-Aldrich) and dexmedetomidine (Patterson Veterinary Supply, Greeley, CO), the peripheral non-selective βAR antagonist nadolol (Sigma-Aldrich), the DBH inhibitor nepicastat (Synosia Therapeutics, Basel, Switzerland), the β1AR antagonist betaxolol (Sigma-Aldrich), the β2AR antagonist ICI 118,551 (Sigma-Aldrich), the α2AR antagonist atipamezole (Patterson Veterinary Supply), the peripheral aromatic acid decarboxylase inhibitor benserazide (Sigma-Aldrich), and the synthetic NE precursor l-3,4-dihydroxyphenylserine (DOPS; Lundbeck, Deerfield, IL). All drugs were dissolved in sterile saline (0.9% NaCl) except for prazosin, which was dissolved in saline containing 1.5% DMSO + 1.5% Cremophor EL, and DOPS, which was dissolved in distilled water with 2% HCl, 2% NaOH, and 2 mg/kg vitamin C.

All compounds except DOPS were administered by i.p. injections at a volume of 10 ml/kg. Sterile saline vehicle was injected to control for any confounding effect of injection stress on behavior, and vehicle-treated animals were used for statistical comparison with anti-adrenergic compounds. For the DOPS rescue experiment, *Dbh -/-* mice were injected with DOPS (1 g/kg, s.c.) + benserazide (250 mg/kg; Sigma-Aldrich), then tested 5 h later once NE levels peaked. The vehicle control for the DOPS experiment was also administered 5 h before testing (Rommelfanger et al. 2007; Schank et al. 2008; Thomas et al. 1998). The doses for all compounds tested were based on previous studies (Durcan et al. 1989; Ji et al. 2014; Kauppila et al. 1991; Luttinger et al. 1985; Millan et al. 2000; Murchison et al. 2004; Rudoy et al. 2007; Schank et al. 2008; Schroeder et al. 2013; Stemmelin et al. 2008; Van Der Laan et al. 1985) and optimized in pilot experiments to ensure that behavioral effects were not due to sedation.

### Elevated zero maze

Mice were exposed to the EZM (2” wide track, 20” diameter) under low light for 5 min (Shepherd et al. 1994; Tillage et al. 2019). Time spent in the open and closed segments of the maze, entries into the open segments, distance traveled, and velocity were recorded on an overhead camera and measured using TopScan software (Clever Sys Inc., Reston, VA).

### Nestlet shredding

Individual mice were removed from their home cages and placed into a new standard mouse cage (13” x 7” x 6”) with clean bedding and a standard cotton nestlet square (5 cm x 5 cm, roughly 3g). The nestlets were pre-weighed before the start of testing to calculate the % shredded at the end of the task (Angoa-Pérez et al. 2013; Angoa-Pérez et al. 2012; Li et al. 2006). Mice were left undisturbed for 30, 60, or 90 min, after which they were returned to their home cages. The weights of the remaining non-shredded nestlet material were recorded, as previously described (Angoa-Pérez et al. 2013; Li et al. 2006). In one experiment, mice were singly housed with nestlets for 24 h. For pharmacological experiments, all drugs except DOPS and nepicastat were administered 15 min before testing, and the task duration was set at 60 min. Nepicastat was administered 2 h prior to testing to allow for maximal NE depletion (Schroeder et al. 2013), and DOPS was administered 5 h before testing (Schank et al. 2008; Thomas et al. 1998). To determine the effect of test cage habituation on NS behavior, individual *Dbh -/-* and control mice were moved into new, clean mouse cages without a nestlet. After 3 h of habituation, a nestlet was introduced to the test cage, and NS within 60 min was measured. The NS behavior of habituated mice was compared with NS behavior in mice tested without habituation.

### Marble burying

Individual mice were removed from their home cages and placed into a large novel cage (10” x 18” x 10”) with 20 black glass marbles arranged in a 4 x 5 grid pattern on top of 2” of normal bedding substrate. The test cage had no lid, and the room was brightly lit. Mice were left undisturbed in the test cage for 30 min. At the end of the test, mice were returned to their home cages. Digital photographs of each test cage were captured at uniform angles and distances. The number of marbles buried was determined by counting the marbles that remained unburied and subtracting this number from 20. A marble was counted as “buried” if at least 2/3 of it was submerged by bedding (Angoa-Pérez et al. 2013; Tillage et al. 2019). For pharmacological experiments, all drugs were administered by i.p. injection 30 min prior to testing, as described (Jimenez-Gomez et al. 2011; Sugimoto et al. 2007).

### c-fos immunohistochemistry

*Dbh -/-* and control mice were exposed to MB or NS tasks, after which they were left undisturbed in the test cage. 90 min following the start of the task, mice were euthanized with an overdose of sodium pentobarbital (Fatal Plus, 150 mg/kg, i.p.; Med-Vet International, Mettawa, IL) and transcardially perfused with cold 4% paraformaldehyde in 0.01 M PBS. After extraction, brains were postfixed for 24 h in 4% paraformaldehyde at 4°C, and then transferred to cryoprotectant 30% sucrose/PBS solution for 72 h at 4°C. Brains were then embedded in OCT medium (Tissue-Tek) and sectioned by cryostat into 40-um coronal slices at the level of the ACC and LC. Another group of *Dbh -/-* and control mice was selected for baseline comparisons of c-fos induction in the LC and ACC; these animals were naïve to the tasks and were removed from their home cages and immediately euthanized and perfused.

Brain sections were blocked for 1 h at room temperature in 5% normal goat serum (NGS) in 0.01 M PBS/0.1% Triton-X permeabilization buffer. Sections were then incubated for 48 h at 4°C in NGS blocking/permeabilization buffer, including primary antibodies raised against c-fos (rabbit anti-c-fos, Millipore, Danvers, MA, ABE457; 1:5000) and NET (mouse anti-NET; MAb Technologies, Neenah, WI, NET05-2; 1:1000). After washing in 0.01 M PBS, sections were incubated for 2 h in blocking buffer, including goat anti-rabbit AlexaFluor 568 and goat anti-mouse AlexaFluor 488 (Invitrogen, Carlsbad, CA; 1:500). After washing, the sections were mounted onto Superfrost Plus slides and coverslipped with Fluoromount-G plus DAPI (Southern Biotech, Birmingham, AL).

### Fluorescent imaging and c-fos quantification

Fluorescent micrographs of immunostained sections were acquired on a Leica DM6000B epifluorescent upright microscope at 20x magnification with uniform exposure parameters. One representative atlas-matched section of the LC and the ACC was selected from each animal for c-fos quantification. An identically sized region of interest was drawn for all images to delineate the borders of both structures in all animals after both behaviors. Image processing and analysis were performed using ImageJ software. Our analysis pipeline includes background subtraction, intensity thresholding (Otsu method), and automated quantification within defined regions of interest, guided by size and shape criteria for c-fos+ cells (size: 50–100 mm^2^, circularity: 0.6–1.0). The NET antibody was used to define LC cell bodies and identify axon terminals in apposition to ACC neurons.

### Statistical analysis

For NS experiments, the effects of time on shredding in *Dbh -/-* and *Dbh +/-* mice were compared using a two-way repeated measures ANOVA (genotype x time), with post hoc Sidak’s test for multiple comparisons. The effects of pre-habituation to the test cage on shredding were also compared by two-way repeated measures ANOVA for each genotype (genotype x habituation time), with post hoc Sidak’s test for multiple comparisons. NS behavior between genotypes after 24 h was compared using an unpaired t-test. NS behavior in DOPS- and vehicle-treated *Dbh -/-* mice was also assessed by unpaired t-test. The effects of anti-adrenergic drugs on NS compared to vehicle were assessed using a one-way ANOVA, with post hoc Dunnett’s test for multiple comparisons. For the MB experiments, genotype differences in burying were assessed by unpaired t-test. The effects of anti-adrenergic drugs on MB compared to vehicle were assessed by one-way repeated-measures ANOVA, with post hoc Dunnett’s test for multiple comparisons.

For c-fos quantification, genotype differences were compared in the LC and ACC at baseline and after MB or NS. Comparisons were made within behavioral tasks and between genotypes by multiple t-tests using the Holm-Sidak correction for multiple comparisons. The threshold for adjusted significance was set at *p* < 0.05, and two-tailed variants of tests were used throughout. Graphical data are presented as group mean ± SEM. Statistical analyses were conducted and graphs were made using Prism Version 7 (GraphPad Software, San Diego, CA).

## Results

### Central norepinephrine is necessary and sufficient for stress-induced nestlet shredding behavior

A cohort of age- and sex-matched *Dbh -/-* and *Dbh +/-* control mice were compared in the NS task, with three different task durations on three different test days: 30 min on Day 1, 60 min on Day 2, and 90 min on Day 3. A two-way repeated measures ANOVA (genotype x task duration) showed a main effect of time (F(2,24) = 20.93, *p* < 0.0001), a main effect of genotype (F(1,12) = 171.2, *p* < 0.0001), and a time x genotype interaction (F(2,24) = 15.81, *p* < 0.001). Post hoc analyses revealed that control mice shredded more nestlets at 60 min (75.44%) and 90 min (96.16%) than at 30 min (22.82%) [*p* < 0.0001], while shredding was minimal in *Dbh -/-* mice at all time points (*p* > 0.05 for all *Dbh -/-* time point comparisons) (Fig. 1a). After animals were individually housed with nestlets for 24 h, all mice of both genotypes shredded 100% of their nestlets (t(14) = 0, *p* > 0.99) (Fig. 1b). Acute restoration of central NE synthesis in *Dbh -/-* mice with DOPS + benserazide increased shredding to control levels at the 60-min time point (81.68% vs 6.88%; t(9) = 10.97, *p* < 0.0001) (Fig. 1c). Finally, the effect of a 3-h habituation to the test cage on NS (60-min time point) was compared between genotypes. A repeated measures two-way ANOVA (genotype x habituation status) showed a main effect of habituation status (F(1,10) = 9.47, *p* = 0.01), a main effect of genotype (F(1,10) = 23.62, *p* < 0.001), and a strong trend for a genotype x habituation status interaction [F(1,10) = 4.71, *p* = 0.06]. Post hoc comparisons revealed that habituation reduced NS in control mice compared to no habituation (74.1% vs 34.84%; *p* < 0.01), but NS in *Dbh -/-* mice did not differ significantly after habituation (14.19% vs 7.41%; *p >* 0.05) (Fig. 1d).

**Fig. 1.**
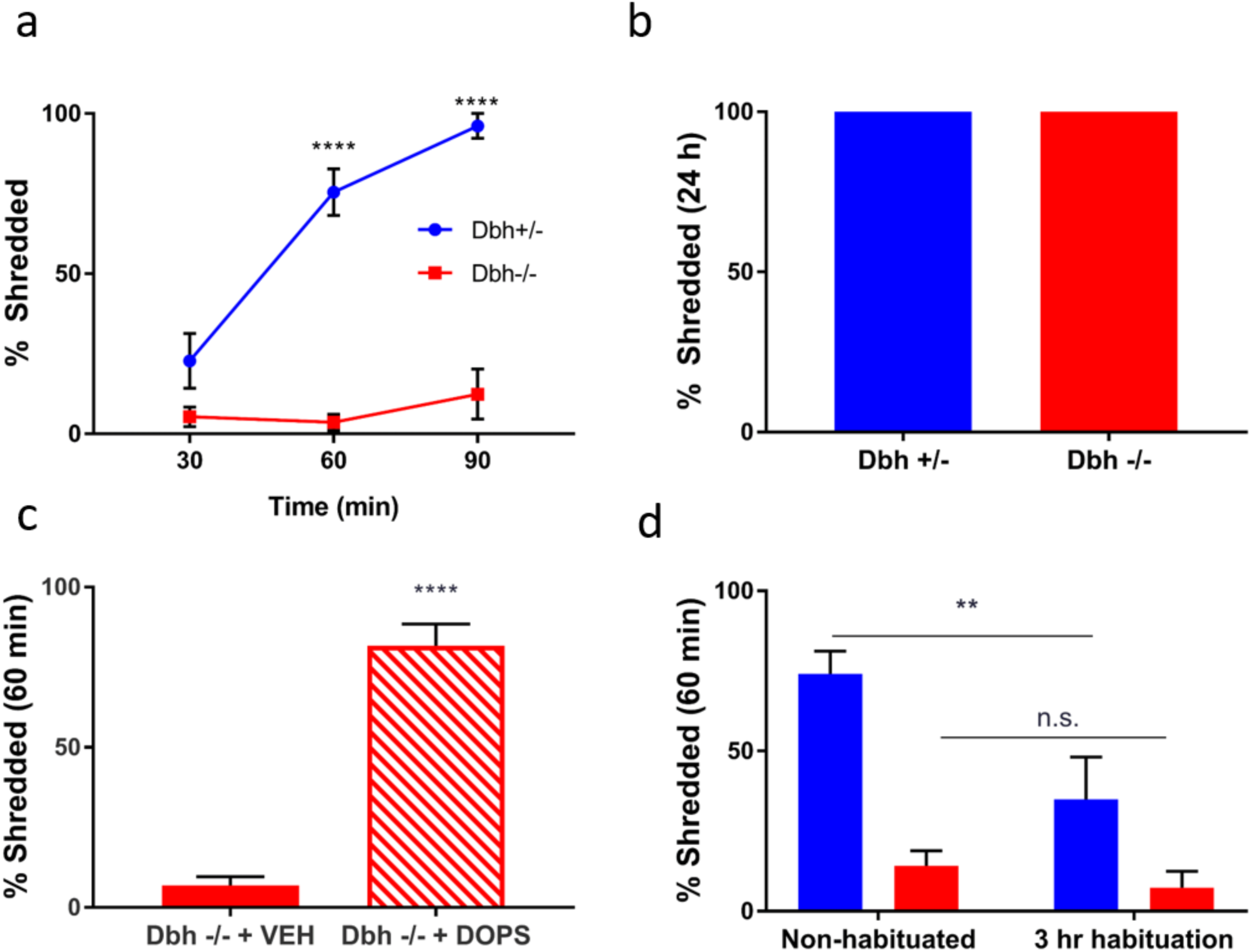
Assessment of nestlet shredding behavior in *Dbh -/-* and *Dbh +/-* mice. **a** Regardless of task duration, NS behavior was profoundly reduced in *Dbh -/-* mice (n = 7) compared to controls (n = 7). In control mice, NS increased when task duration was extended to 60 or 90 min compared to 30 min, but NS behavior in *Dbh-/-* mice (n = 7) did not increase as a function of task duration. **b** When mice were singly housed with nestlets overnight, NS reached 100% and did not differ between *Dbh -/-* (n = 8) and control mice (n = 8). **c** Restoring central NE levels in *Dbh -/-* mice by treating them with DOPS (1 g/kg) + benserazide (250 mg/kg) 5 h prior to testing (n = 5) increased NS compared to vehicle (n = 6). **d** 3-h habituation to the test cage significantly reduced NS in control mice (n = 6), indicating that shredding was mediated by acute cage-change stress. Habituation did not affect NS in *Dbh -/-* mice (n = 6). \*\*\*\**p* < 0.0001, **p *<* 0.01, n.s. = not significant.

### Nestlet shredding can be suppressed in control mice by anti-adrenergic drugs

Because *Dbh +/-* mice demonstrated robust but not maximal NS behavior at 60 min (Fig. 1a,d), we used this task duration for pharmacological experiments. Control mice were administered either saline vehicle or experimental compounds in their home cages 15 min prior to testing. The test period began once the mouse was placed into a new clean cage with a pre-weighed cotton nestlet square. The experimental compounds used here were the following anti-adrenergic drugs and their mechanisms of action: the α1AR antagonist prazosin (0.5 mg/kg), the βAR antagonist propranolol (10 mg/kg), the α2AR antagonist atipamezole (0.5 mg/kg), the α2AR agonists guanfacine (0.3 mg/kg) and dexmedetomidine (0.02 mg/kg), and the DBH inhibitor nepicastat (100 mg/kg). A one-way ANOVA showed a main effect of treatment (F(6,41) = 126.3, *p* < 0.0001). Post hoc comparisons revealed that propranolol (*p* < 0.0001), guanfacine (*p* < 0.0001), dexmedetomidine (*p* < 0.0001), and nepicastat (*p* < 0.0001) reduced NS compared to vehicle. There was no significant difference in NS in the prazosin group compared to saline (*p* > 0.05). By contrast, atipamezole significantly increased NS behavior compared to vehicle (*p =* 0.03) (Fig. 2a).

**Fig. 2.**
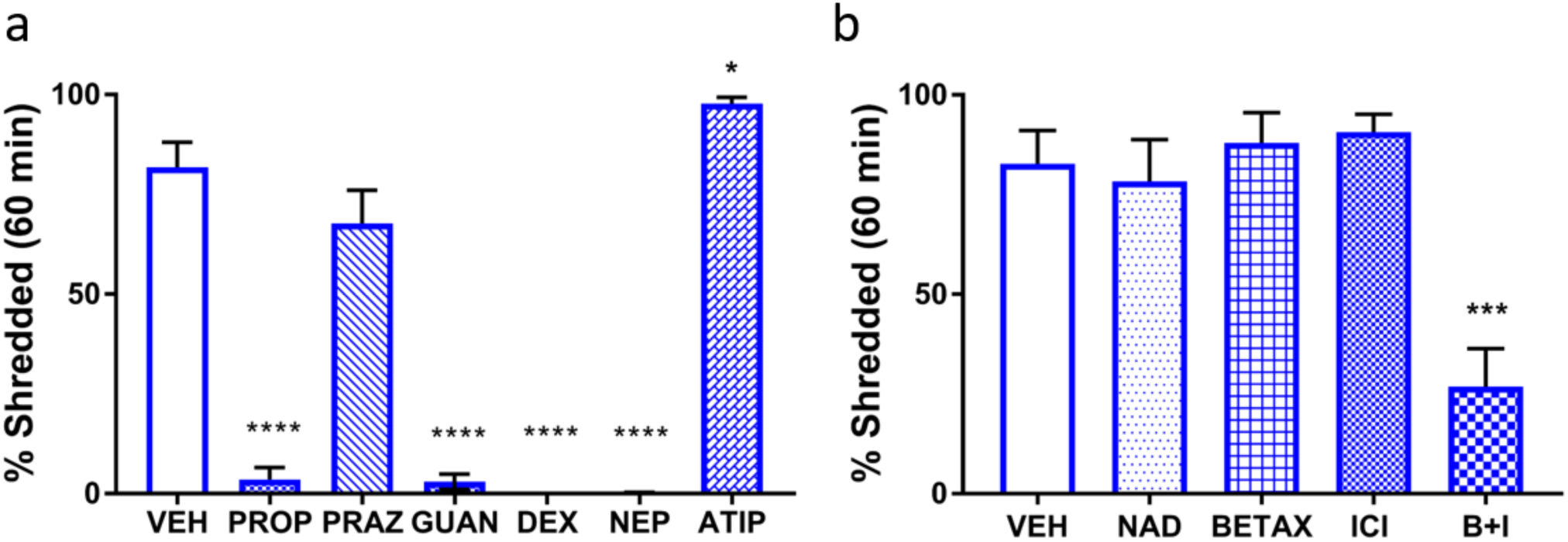
Anti-adrenergic drugs suppress nestlet shredding in control mice. **a** Compared to vehicle-treated mice (VEH; n = 7), NS was reduced by propranolol (10 mg/kg; PROP; n = 7), guanfacine (0.3 mg/kg; GUAN; n = 7), dexmedetomidine (0.02 mg/kg; DEX; n = 7), and nepicastat (100 mg/kg, NEP; n = 7). NS was enhanced by atipamezole (0.5 mg/kg; ATIP; n = 7) and unaffected by prazosin (0.5 mg/kg; PRAZ; n = 6). **b** Compared to VEH-treated mice (n = 6), nadolol (10 mg/kg; NAD; n = 6), betaxolol (5 mg/kg; BETAX; n = 7), and ICI-118,551 (1 mg/kg; ICI; n = 7) were ineffective at reducing NS, but the combination of BETAX and ICI (B+I; n = 7) reduced NS compared to VEH. \*\*\**p* < 0.001.

To rule out the contribution of peripheral βARs and determine whether different subtypes of central βAR regulate the expression of NS, another group of *Dbh +/-* mice was tested in a separate series of experiments. A one-way ANOVA of treatment efficacy showed a main effect of treatment (F(4, 28) = 10.91, *p* < 0.0001). Post hoc analyses revealed that the peripheral βAR antagonist nadolol (10 mg/kg) was ineffective at reducing NS compared to vehicle (p > 0.05), as was the selective β1 antagonist betaxolol (5 mg/kg) (*p* > 0.05) and the selective β2 antagonist ICI 118,551 (1 mg/kg) (*p* > 0.05). However, a cocktail of the β1 and β2AR antagonists was sufficient to reduce NS compared to vehicle (*p <* 0.001), consistent with a partially redundant role for β1 and β2 ARs in NS behavior (Fig. 2b).

### Genetic or pharmacological suppression of norepinephrine transmission attenuates marble burying

*Dbh -/-* and *Dbh +/-* control mice were assessed in the MB task. Compared to control mice, *Dbh -/-* mice buried fewer marbles (t(26) = 2.37, *p* = 0.03) (Fig. 3a). Because NE is important for arousal and wakefulness, a subset of mice was filmed to determine whether *Dbh* -/- mice were not burying marbles because they were falling asleep. Neither control nor *Dbh -/-* mice fell asleep during the MB task, and mice of both genotypes engaged in typical behaviors, including grooming, rearing, and sniffing. We did notice that some *Dbh -/-* mice occasionally exhibited qualitatively unusual behaviors not observed in control mice, such as sitting on top of the marbles (data not shown).

**Fig. 3.**
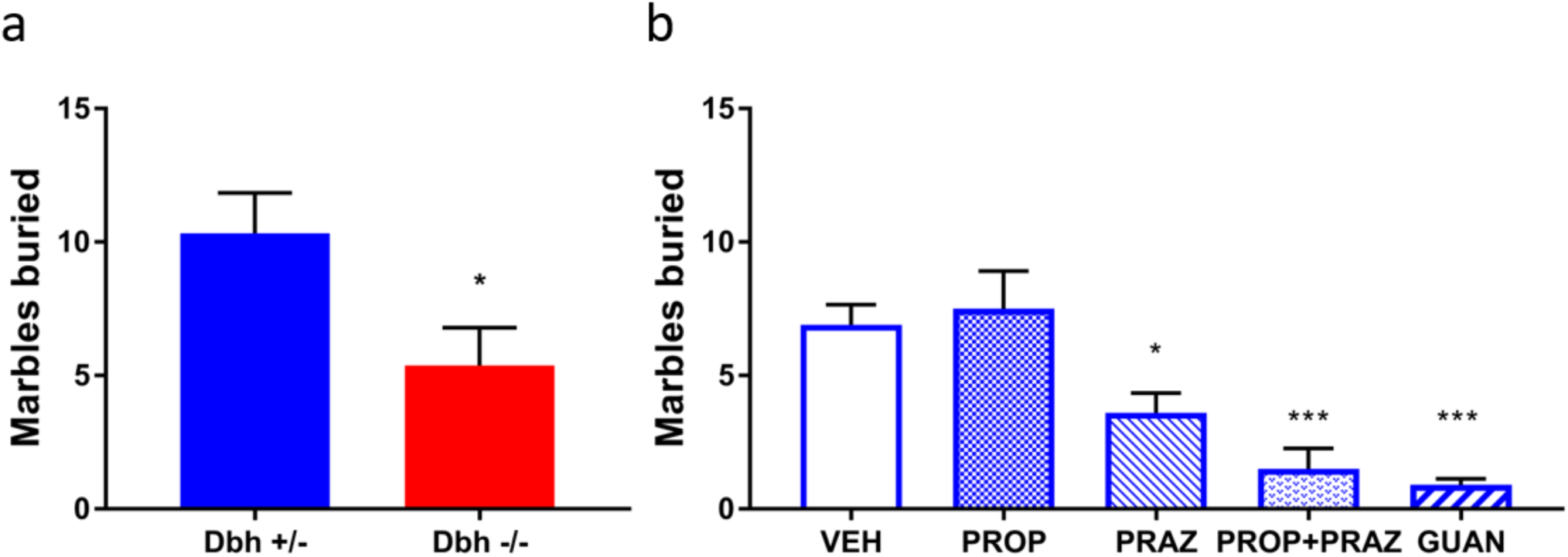
Consequences of genetic and pharmacological disruption of norepinephrine transmission on marble burying. **a** Compared to *Dbh* +/- controls (n = 15), *Dbh -/-* (n = 13) mice buried fewer marbles. **b** In control mice, PROP (10 mg/kg) was ineffective, but PRAZ (0.5 mg/kg), co-administration of PRAZ + PROP, or GUAN (0.3 mg/kg) reduced MB compared to VEH (n = 10, repeated measures). \**p* < 0.05, \*\*\**p* < 0.001.

*Dbh +/-* control mice were used for the pharmacological characterization of anti-adrenergic drug effects on MB. The experimental compounds used here were prazosin (0.5 mg/kg), propranolol (10 mg/kg), a cocktail of prazosin and propranolol at the same doses, and guanfacine (0.3 mg/kg). A one-way repeated measures ANOVA of treatment efficacy revealed a main effect of treatment (F(2,18) = 12.11, *p* < 0.001). Post hoc comparisons demonstrated that, compared to vehicle, prazosin (*p* < 0.05), prazosin + propranolol (*p* < 0.001), and guanfacine (*p* < 0.001) potently suppressed MB compared to vehicle, while propranolol alone was not effective (*p* > 0.05) (Fig. 3b).

### Dbh -/- mice exhibit normal anxiety-like behavior in the elevated zero maze

*Dbh -/- and Dbh +/-* mice have previously been compared in conflict-based anxiety tasks, including the elevated plus maze (EPM), open field test, and light/dark box, and no knockout phenotypes have been observed (Marino et al. 2005; Schank et al. 2008). To confirm that their MB and NS phenotypes were unique to stress-induced repetitive behaviors and unrelated to general anxiety, we evaluated the performance of knockouts and controls in the EZM, another canonical anxiety assay. There were no statistically significant genotype differences on anxiety-like measures in this task, including the percent of time spent in the open segment (t(17) = 0.01, *p* > 0.05) (Fig. 4a) or open segment entries (t(17) = 1.10, *p* > 0.05) (Fig. 4b). *Dbh -/-* mice did exhibit diminished locomotor activity during the EZM task; they traveled at a reduced average velocity compared to controls (t(17)=2.34, *p* = 0.03) (Fig. 4c) and tended to travel shorter distances, although the latter did not reach statistical significance (t(17) = 1.92, *p* = 0.07) (Fig. 4d). Attenuation of these locomotor measures of arousal in *Dbh -/-* mice is consistent with previous reports that these animals exhibit attenuated locomotor responses to novel environments (Porter-Stransky et al. 2019; Weinshenker et al. 2002).

**Fig. 4.**
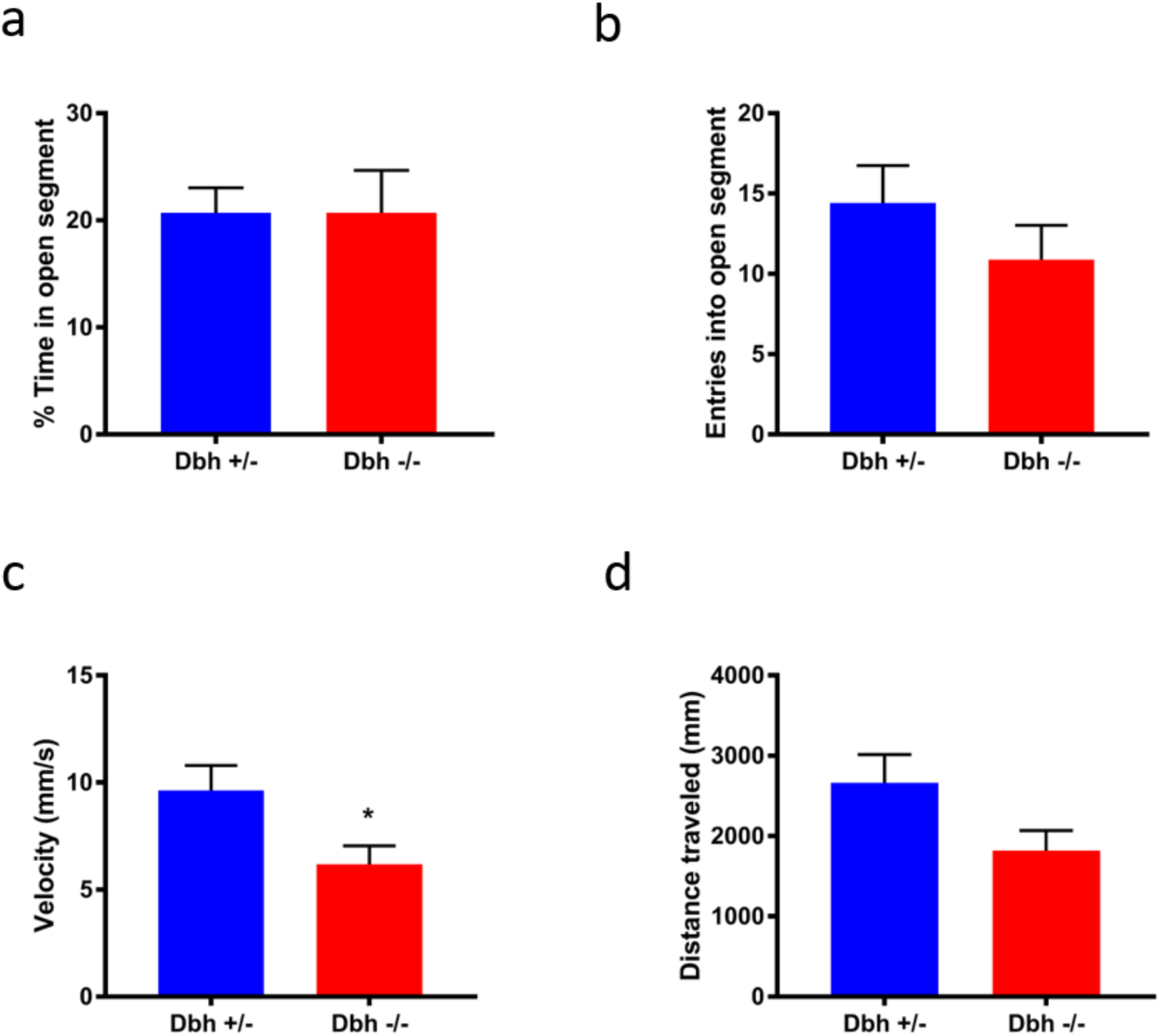
Assessment of *Dbh -/-* and *Dbh +/-* mice in the elevated zero maze. *Dbh -/-* mice did not differ from controls on anxiety-like measures, including **a** % time in the open (anxiogenic) segment and **b** entries into the open segment. There was a significant genotype difference in **c** average velocity and a trend for **d** total distance traveled. N = 9-10 per group, \**p < 0.05*.

### NE-deficient mice show diminished marble burying- and nestlet shredding-induced c-fos expression in anterior cingulate cortex

*Dbh -/-* and *Dbh +/-* control mice were euthanized for quantification of c-fos+ cells in the LC and ACC at baseline and following either NS or MB. At baseline, c-fos expression was minimal and there were no significant genotype differences in c-fos+ cells in either the LC (t(4) = 0.76, *p* > 0.05) or the ACC (t(4)=0.39, *p* > 0.05) (Fig. 5a). Both MB and NS induced robust c-fos expression in both regions, but genotype differences emerged. Following MB, *Dbh -/-* mice had significantly fewer c-fos+ cells in the ACC (t(7) = 6.38, *p* < 0.001), but normal c-fos+ induction in the LC (t(7) = 0.07, *p* > 0.05) (Fig. 5b). Following NS, *Dbh -/-* mice had significantly fewer c-fos+ cells in the ACC (t(10) = 5.77, *p* < 0.001) and the LC (t(10) = 5.66, *p* < 0.001) compared to control mice (Fig. 5c).

**Fig. 5.**
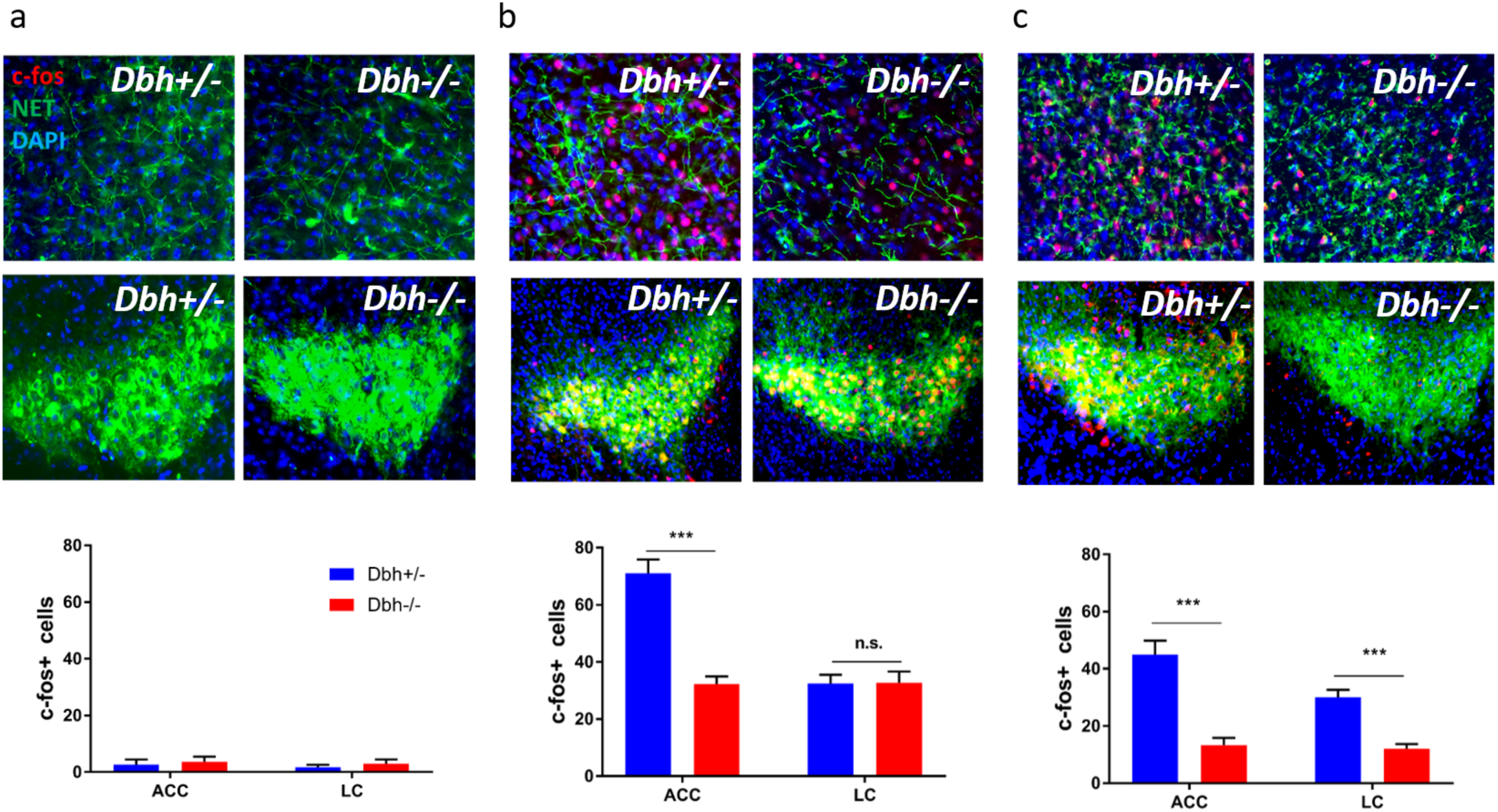
Comparison of nestlet shredding- and mable burying-induced c-fos expression in anterior cingulate cortex and locus coeruleus between *Dbh -/-* and *Dbh +/-* mice. **a** In experimentally naïve mice at baseline, c-fos induction was minimal in the ACC (top row) and LC (bottom row), and there were no differences in c-fos+ cells between *Dbh -/-* (n = 3) and control mice (n = 3) in either region. **b** After MB, fewer c-fos+ cells were detected in the ACC of *Dbh* -/- mice (n = 4) compared to control mice (n = 5), but no genotype differences were found in the LC. **c** After NS, fewer c-fos+ cells were detected in both the ACC and LC of *Dbh* -/- mice (n = 6) compared to control mice (n = 6). \*\*\**p* < 0.001, n.s. = not significant.

## Discussion

### Central norepinephrine is necessary and sufficient for stress-induced nestlet shredding behavior

In this study, we compared NS behavior induced by cage-change stress in NE-deficient and control mice. We found that over the course of 90 min, NE-deficient mice demonstrated virtually no NS, whereas their NE-competent littermates vigorously shredded nearly 100% of their nestlets. Importantly, we observed no genotype differences in NS behavior when mice were given 24 h to shred, indicating that the NE-deficient mice are capable of shredding but simply do not do so with any urgency when placed in a new cage. This finding provides evidence that NE is necessary for rapid NS behavior following cage change, but not for typical nest-building behavior in general. When we acutely restored central NE to *Dbh -/-* mice using DOPS + benserazide, rapid NS behavior was rescued to control levels. This key finding demonstrates that transient pharmacological restoration of central NE synthesis and transmission is sufficient to support stress-induced NS behavior in NE-deficient animals.

### Norepinephrine-deficient mice bury fewer marbles in the marble burying task

Although MB is often interpreted as an anxiety-like behavior, performance in the MB task does not correlate with behavioral measures in other models of anxiety, such as the EPM or open field test (Jimenez-Gomez et al. 2011; Thomas et al. 2009). However, MB behavior is highly correlated with NS behavior and does not habituate after repeated exposures, indicating that it better reflects compulsive behavior (Angoa-Pérez et al. 2013; Li et al. 2006; Witkin 2008). Indeed, NE-deficient mice exhibited normal anxiety-like behavior in the EZM but buried fewer marbles than controls in the MB task. This dissociation in behavioral phenotypes is consistent with previously published reports of normal *Dbh -/-* behavior in canonical conflict-based anxiety paradigms (Marino et al. 2005; Schank et al. 2008) and suggests that NE may play a more important role in the expression of stress-induced compulsive behaviors than in innate anxiety (Brady 1994; Kedia and Chattarji 2014; Mantsch et al. 2010; McCall et al. 2015; Valentino et al. 1993).

### Nestlet shredding and marble burying can be suppressed in control mice by anti-adrenergic drugs

NE acts primarily through α1 and βARs to exert neuromodulatory effects on target cells, and NE transmission can be decreased by activation of α2 inhibitory autoreceptors. To determine which ARs are involved in the expression of stress-induced NS behavior, we treated control mice with a battery of anti-adrenergic drugs that block ARs on target cells (α1 and βAR antagonists), activate inhibitory autoreceptors (α2AR agonists), or prevent NE biosynthesis (DBH inhibitors). Prazosin, an α1AR antagonist, did not significantly affect NS compared to saline vehicle. However, disruption of NE synthesis with nepicastat, reduction of NE release with the α2AR agonists guanfacine and dexmedetomidine, or antagonism of βARs with propranolol profoundly suppressed NS behavior in control mice. In a separate set of experiments to determine the requirement of central β1 and β2 AR activation for NS behavior, we found that blockade of βARs outside the brain with the peripherally restricted antagonist nadolol did not suppress NS. We also found that selective antagonism of either β1ARs or β2ARs alone was not sufficient to suppress NS, but NS could be reduced by co-administration of β1- and β2AR-selective antagonists. Thus, the effect of propranolol on NS behavior is likely to be mediated by central β1 and β2ARs, which have overlapping distributions in some brain regions like the ACC, amygdala, and hippocampus, and may serve partially redundant signaling functions (Abraham et al. 2008; Qu et al. 2008; Rainbow et al. 1984; Zheng et al. 2015; Zhou et al. 2013).

NS behavior was increased when NE transmission was enhanced via the α2AR antagonist atipamezole, which blocks α2 inhibitory autoreceptors. In addition, we found that anti-adrenergic drugs potently suppressed MB behavior in control mice. Guanfacine and prazosin, but not propranolol, dramatically reduced MB. These results are consistent with the literature implicating α1AR activation in behavioral reactivity to novelty (Stone et al. 2006), maintenance of generalized arousal (Broese et al. 2012; Porter-Stransky et al. 2019; Stone et al. 2003), and stress-induced anxiety (Rasmussen et al. 2016; Skelly et al. 2014).

### NE-deficient mice show diminished c-fos induction in anterior cingulate cortex following nestlet shredding and marble burying

The ACC is implicated in the pathophysiology of OCD (Brennan et al. 2015; McGovern and Sheth 2017), and surgical ablation of the ACC can relieve OCD symptoms (Jung et al. 2006; Kim et al. 2003). Furthermore, the ACC expresses all subtypes of ARs (Crino et al. 1993; Rainbow et al. 1984) and is bidirectionally connected to the LC (Gompf et al. 2010; Loughlin et al. 1982). Baseline genotype differences in c-fos induction in the LC and ACC were not seen when animals were euthanized immediately after removal from their home cages. However, c-fos induction was dramatically reduced in the ACC of NE-deficient mice after NS and MB compared to controls. After NS, but not MB, c-fos induction was reduced in the LC of NE-deficient mice compared to controls.

The reason for the emergence of genotype differences in LC activity following NS but not MB is unclear but could be related to task duration and complexity, or possibly to engagement of other structures and neurotransmitter systems that were not considered in our analysis. Given that hyperactivity of the ACC (Fitzgerald et al. 2005; Mavrogiorgou et al. 2002), reduced endocrine response to clonidine (Hollander et al. 1991; Siever et al. 1983), and abnormal catecholamine metabolism (Benkelfat et al. 1991; Schindler et al. 2000) have all been reported in patients with OCD, we propose that excessive NE transmission to the ACC may account for some of the cognitive and affective symptoms of OCD (De Geus et al. 2007).

### Clinical implications

Although clinical studies examining the effects of α1AR blockade on OCD symptoms are scarce (Feenstra et al. 2016), our findings suggest that further clinical trials with prazosin for OCD patients are justified. The ability of guanfacine to suppress MB is consistent with evidence from the limited number of studies showing that other α2AR agonists like clonidine and dexmedetomidine can reduce MB (Millan et al. 2000; Young et al. 2006). However, clonidine and dexmedetomidine have potent sedative properties, which can hamper the interpretation of their behavioral effects in rodents and constrains their tolerability for psychiatric patients. At the dose used in this study, guanfacine had no sedative or motor-impairing effects; in fact, the sedative and motor-impairing ED_50_ for guanfacine in rodents is roughly five-fold higher than the dose used in the present study (Luttinger et al. 1985; Scholtysik 1980; Van Der Laan et al. 1985). Clinically, guanfacine is preferred over clonidine for treatment of pediatric ADHD due to its lack of sedative effects and superior tolerability (Arnsten et al. 1988; Posey and McDougle 2007; Sallee and Eaton 2010; Sallee et al. 2009). Propranolol and guanfacine are well tolerated by patients and have been used both on- and off-label for the treatment of other psychiatric disorders (Chappell et al. 1995; Cummings et al. 2002; Lederman 1999; Steenen et al. 2016). Nepicastat and guanfacine are effective for attenuating compulsive drug-seeking behavior in animal models of addiction (Colombo et al. 2014; Le et al. 2011; Schroeder et al. 2013), as well as relapse in patients with substance abuse disorders (De La Garza II et al. 2015). In scattered psychiatric case studies, the α2AR agonists guanfacine and clonidine have substantially improved OCD symptoms in certain patients (Knesevich 1982; Lipsedge and Prothero 1987; Taormina et al. 2016).

### Limitations and future directions

It is noteworthy that 5-HT-deficient and NE-deficient mice have diametrically opposite phenotypes in the MB and NS tasks (Angoa-Pérez et al. 2013; Angoa-Pérez et al. 2012). Disentangling the role of the NE and 5-HT neuromodulatory systems in the expression of stress-induced compulsive behaviors will be a challenging but worthwhile endeavor. From a circuitry standpoint, it is generally agreed that serotonergic neurons in the raphe nuclei exert a tonic inhibitory influence on the LC, while the LC exerts a tonic excitatory influence on serotonergic neurons in the dorsal raphe (Kim et al. 2004; Pudovkina et al. 2002; Segal 1979; Szabo and Blier 2001). Thus, a possible explanation for the excessive MB and NS behavior observed in 5-HT-deficient mice is that the LC may be hyperactive in the absence of serotonergic modulation. This hypothesis could be tested by administering anti-adrenergic agents, such as nepicastat or guanfacine, to 5-HT-deficient mice and measuring NS and MB behavior when NE transmission is reduced.

A limitation of our study is that we cannot infer a causal role for LC-NE transmission to the ACC in the expression of NS and MB behaviors. More sophisticated studies using tools to manipulate distinct circuits will be required to functionally dissect the NE-dependent network that governs OCD-like behavior. Furthermore, our finding that βARs mediate NS, while α1ARs mediated MB, suggests that different cell types and/or circuits may support these correlated but distinct behaviors. The requirement of α1AR activation for MB but not NS may be related to the size and complexity of the test cage for each task. For MB, testing occurred in large cages “enriched” with marbles, whereas NS was performed in a normal mouse cage with clean bedding and a standard nestlet square. The relative complexity of the MB test environment might activate arousal-promoting α1ARs that are not engaged in the simpler NS environment (Stone et al. 1999; Stone et al. 2005; Stone et al. 2004; Stone et al. 2006). The requirement of βARs for NS but not MB may be related to the longer duration of the task (60 min for NS vs 30 min for MB), which could allow for synergistic actions of corticosterone on βAR signaling in stress-responsive brain regions (Gorman et al. 1993; Roozendaal et al. 2004; Roozendaal et al. 2006).

Finally, this study focused on neuronal activity in two brain regions, the LC and ACC, following MB and NS. The central NE system is anatomically and functionally complex (Robertson et al. 2013; Robertson et al. 2016; Uematsu et al. 2015), and discrete manipulations of genetically defined subpopulations of NE neurons can produce unique or even opposite effects on behavior (Aston-Jones and Waterhouse 2016; Chen et al. 2019; Flavin and Winder 2013; McCall et al. 2017). Moving forward, it will be important to consider the contribution of noradrenergic brainstem groups besides the LC, such as the A2 neurons residing in the nucleus of the solitary tract, to the expression of these behaviors (Itoi and Sugimoto 2010; Rinaman 2010).

### Conclusions

In summary, these findings support an expanded model of OCD pathophysiology that incorporates dysregulation of central NE signaling, possibly between the LC and ACC. In addition, we propose that anti-adrenergic agents should be assessed for clinical efficacy in treating OCD, since many of these drugs are already used both on- and off-label for the treatment of related psychiatric diseases. Given the high cost and failure rate of psychiatric drug testing, the possibility of repurposing an approved drug such as guanfacine to treat OCD represents an attractive alternative to new drug development (Lee et al. 2016).

## Acknowledgments

We thank Lundbeck for providing the DOPS, Synosia Therapeutics for providing the nepicastat, and C. Strauss for helpful editing of the manuscript. This work was supported by the National Institutes of Health (AG061175, NS102306, and DA038453 to DW; GM8602-22 to DL; MH116622 to RPT).

## Conflicts of Interest

DW is co-inventor on a patent concerning the use of selective dopamine β-hydroxylase inhibitors for treatment of cocaine dependence (US-2010-0105748-A1; “Methods and Compositions for Treatment of Drug Addiction”). The other authors declare no conflicts of interest.

